# Increased aerosol transmission for B.1.1.7 (alpha variant) over lineage A variant of SARS-CoV-2

**DOI:** 10.1101/2021.07.26.453518

**Authors:** Julia R. Port, Claude Kwe Yinda, Victoria A. Avanzato, Jonathan E. Schulz, Myndi G. Holbrook, Neeltje van Doremalen, Carl Shaia, Robert J. Fischer, Vincent J. Munster

## Abstract

Airborne transmission, a term combining both large droplet and aerosol transmission, is thought to be the main transmission route of SARS-CoV-2. Here we investigated the relative efficiency of aerosol transmission of two variants of SARS-CoV-2, B.1.1.7 (alpha) and lineage A, in the Syrian hamster. A novel transmission caging setup was designed and validated, which allowed the assessment of transmission efficiency at various distances. At 2 meters distance, only particles <5 µm traversed between cages. In this setup, aerosol transmission was confirmed in 8 out of 8 (N = 4 for each variant) sentinels after 24 hours of exposure as demonstrated by respiratory shedding and seroconversion. Successful transmission occurred even when exposure time was limited to one hour, highlighting the efficiency of this transmission route. Interestingly, the B.1.1.7 variant outcompeted the lineage A variant in an airborne transmission chain after mixed infection of donors. Combined, this data indicates that the infectious dose of B.1.1.7 required for successful transmission may be lower than that of lineage A virus. The experimental proof for true aerosol transmission and the increase in the aerosol transmission potential of B.1.1.7 underscore the continuous need for assessment of novel variants and the development or preemptive transmission mitigation strategies.

## Introduction

More than one year has passed since the declaration of the severe acute respiratory syndrome coronavirus 2 (SARS-CoV-2) pandemic in March 2020 by the World Health Organization (WHO). Epidemiological data suggests that the principal mode of infection with SARS-CoV-2 is via airborne transmission ^1–5^. The general definition held by the WHO states that large droplets disperse over a short distance and settle in the upper respiratory tract, while aerosols (<5 µm) can form droplet nuclei, travel over long distance, and significantly deposit in the lower respiratory tract ^6^. Both forms are typically considered airborne transmission. For influenza A virus, another respiratory virus, studies have elucidated the airborne potential and discussed the relative contribution of droplets vs. aerosols and the site of viral exposure and shedding ^7–10^. Similar data for SARS-CoV-2 is currently unavailable.

Genetic variants of SARS-CoV-2 continue to be detected worldwide and variants of concern (VOCs) are defined by phenotypic changes including enhanced transmission ^11,12^. Transmissibility is a function of infectiousness, susceptibility, contact patterns between individuals, and environmental stress on the pathogen during transmission ^13^. In our previous work, we have demonstrated that aerosol exposure increases severity of disease in the Syrian hamster and that airborne transmission with a lineage A variant over short distances (<10 cm) is very efficient ^14,15^. However, in these studies we could not differentiate between large particles and true aerosols. No study so far has demonstrated the potential of SARS-CoV-2 for true aerosol transmission with particles <5 µm. Here, we specifically designed transmission cages to model aerosol transmission over 2 meters distance, at which only particles <5 µm traverse. We showed highly efficient aerosol transmission of SARS-CoV-2 at 2 meters distance within one hour of exposure. Lastly, we demonstrated increased airborne transmission competitiveness of B.1.1.7 over a lineage A variant.

## Results

### Design and validation of hamster aerosol transmission cages

Direct contact and airborne transmission have been demonstrated in the Syrian hamster model for SARS-CoV-2 ^14,16^. However, demonstration of true aerosol transmission of SARS-CoV-2 should only include particles <5 µm, over longer distances and in the absence of any other potential transmission routes such as fomite or direct contact. To determine if SARS-CoV-2 can transmit successfully via aerosols, we designed and validated a caging system to study the relationship between particle size and distance. The design consisted of two rodent cages connected via a polyvinylchloride (PVC) connection tube (76 mm inside diameter) which allowed airflow, but no direct animal contact, from the donor to the sentinel cage. The distance between donor and sentinel cage could be varied (16.5, 106, or 200 cm) by exchanging the PVC connection tube (**Supplemental Figure 1 A, B**). Directional airflow from the donor to the sentinel cage was generated by negative pressure. The air velocity generated by the airflow through the connection tube averaged at 327, 370, and 420 cm/min for the 16.5, 106, and 200 cm distances, respectively (**Supplementary Table 1**). This allowed for 30 cage changes per hour.

We next validated the caging design using an aerodynamic particle sizer to analyze the aerodynamic size of particles (dynamic range from <0.5-20 µm) traversing from donor to sentinel cage. Droplets and aerosols were generated in the donor cage (20% (v/v) glycerol solution, sprayed with a standard spray bottle) and the particle size profile was determined at the beginning and end of the connecting tube to study the potential for size exclusion of the respective cage setups. The reduction of particles was size and distance dependent. At a distance of 16.5 cm between cages, relatively limited size exclusion of the generated particles was observed; ≥6.9% of particles 5-10 µm and ≥42.8% of particles ≥10 µm did not travel into the sentinel cage (**Fig 1 A/D**). At the intermediate distance of 106 cm between cages an increased reduction of number of particles and size exclusion was observed; ≥70% of particles ≥5 µm did not traverse into the sentinel cage and no particles ≥10 µm were detected. Hence, while in the donor cage 4.86% of detected particles were >5 µm, in comparison the particle profile in the sentinel cage contained only 2% particles >5 µm (**Fig 1 B/E**). At the longest distance of 200 cm, we observed an almost complete size exclusion of particles ≥5 µm; ≥95% of particles 5-10 µm did not traverse and no particle ≥10 µm were detected in the sentinel cage. The composition profile of particles in the sentinel cage comprised only 0.5% particles ≥5 µm (**Fig 1 C/F**). These combined results demonstrate that we have developed a novel caging system to effectively investigate the impact of distance and particle size exclusion on the transmission of SARS-CoV-2. The overall absence of particles ≥10 µm and extensive reduction of particles 5-10 µm indicate that the caging system with the distance of 200 cm is suitable to study true aerosol transmission; whereas, the 16.5 and 106 cm set-ups are suitable to study airborne transmission occurring via droplet, aerosols or a combination thereof.

**Figure 1:**
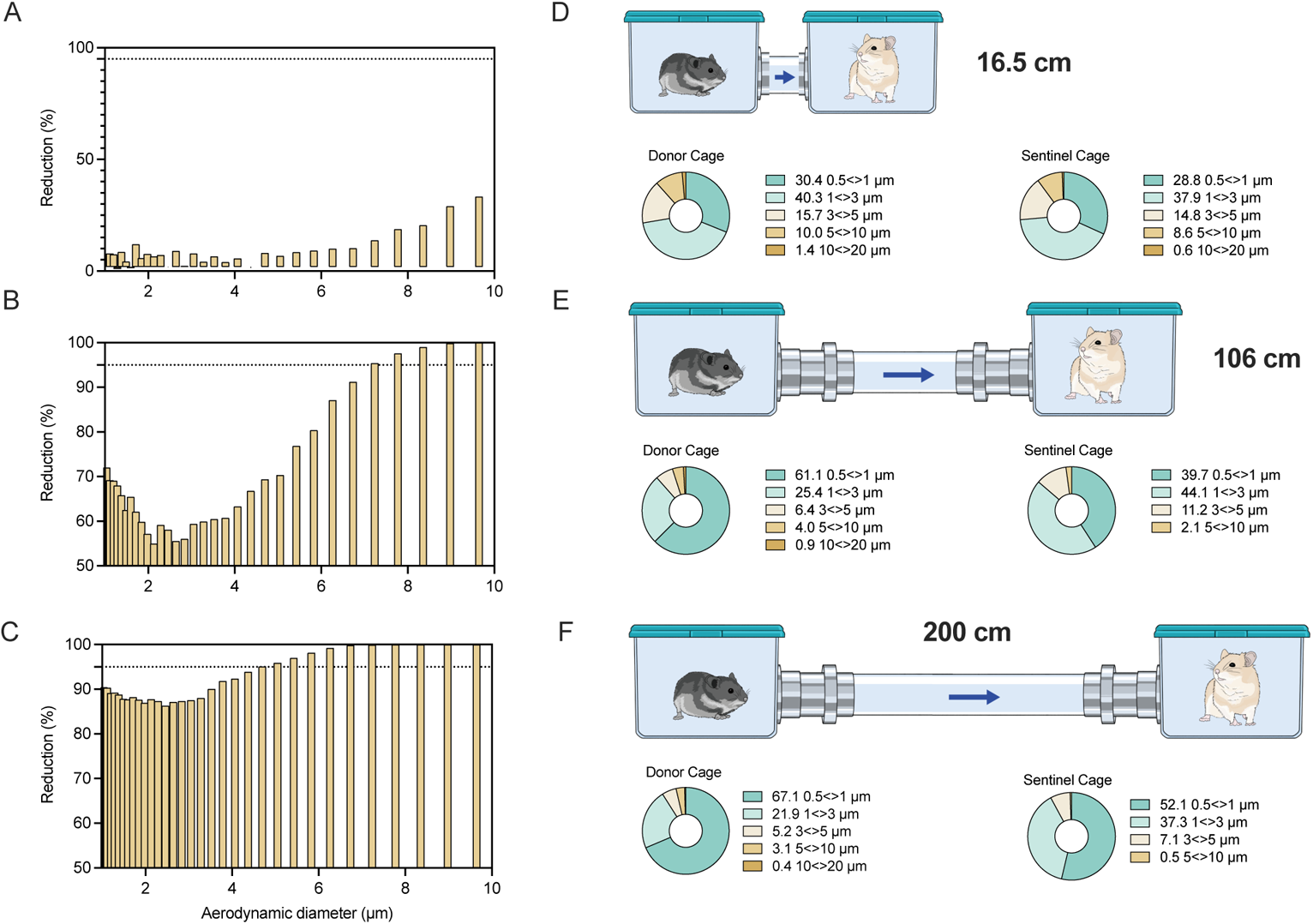
Design and validation of aerosol transmission cages. Transmission cages were designed to model airborne transmission between Syrian hamsters at 16.5 cm, 106 cm and 200 cm distance. Droplets were generated by spraying a 20% glycerol/water solution into the donor cage. Size of particles travelling between donor and sentinel cages were determined. **A/B/C**. Particle reduction by aerodynamic diameter between the donor and sentinel cage at 16.5 cm (A) 106 cm (B) and 200 cm distance (C). Dotted line = 95% reduction in particles. Aerodynamic diameter 1-10 μm. **D/E/F.** Schematic visualization of the transmission cages at 16.5 cm (A), 106 cm (B) and 200 cm distance (C) and corresponding particle distribution detected in each donor and sentinel cage.

### SARS-CoV-2 aerosol transmission over 2 meters distance

Experimental SARS-CoV-2 airborne transmission has been demonstrated over short distances in the Syrian hamster model ^14,15^. Using the validated caging system, we first investigated short-distance airborne transmission. For each distance, four donor animals were inoculated intranasal (I.N) with 8×10^4^ TCID_50_ SARS-CoV-2 lineage A. After 12 hours, the infected animals were placed into the donor (upstream) side of the cages and four sentinels were placed into the downstream cages (2:2 ratio) and were exposed for 72 hours.

At a distance of 16.5 cm, SARS-CoV-2 successfully transmitted to all the sentinels at 12 hours post exposure (HPE) (**Figure 2 A/B**). Genomic (g)RNA and subgenomic (sg)RNA, a marker for replicating virus, were found in oropharyngeal swabs of all sentinels at 48 HPE. At a distance of 106 cm, SARS-CoV-2 gRNA and sgRNA were detected in oropharyngeal swabs of one sentinel as early as 12 HPE. At 48 HPE all sentinels were positive for gRNA and sgRNA in oropharyngeal swabs (**Figure 2 C/D**). At a distance of 200 cm, no respiratory shedding was detectable in any sentinel 12 HPE, but at 48 HPE all sentinels were positive for gRNA and sgRNA in oropharyngeal swabs (**Figure 2 E/F**). Additionally, at 14 days post exposure (DPE) all sentinels from all three groups had seroconverted, as demonstrated by high antibody titers against SARS-CoV-2, measured by anti-spike ELISA (**Supplemental Table 2**). These data demonstrate the ability of SARS-CoV-2 to transmit over long and short distances and suggest that transmission efficiency may be distance dependent.

**Figure 2:**
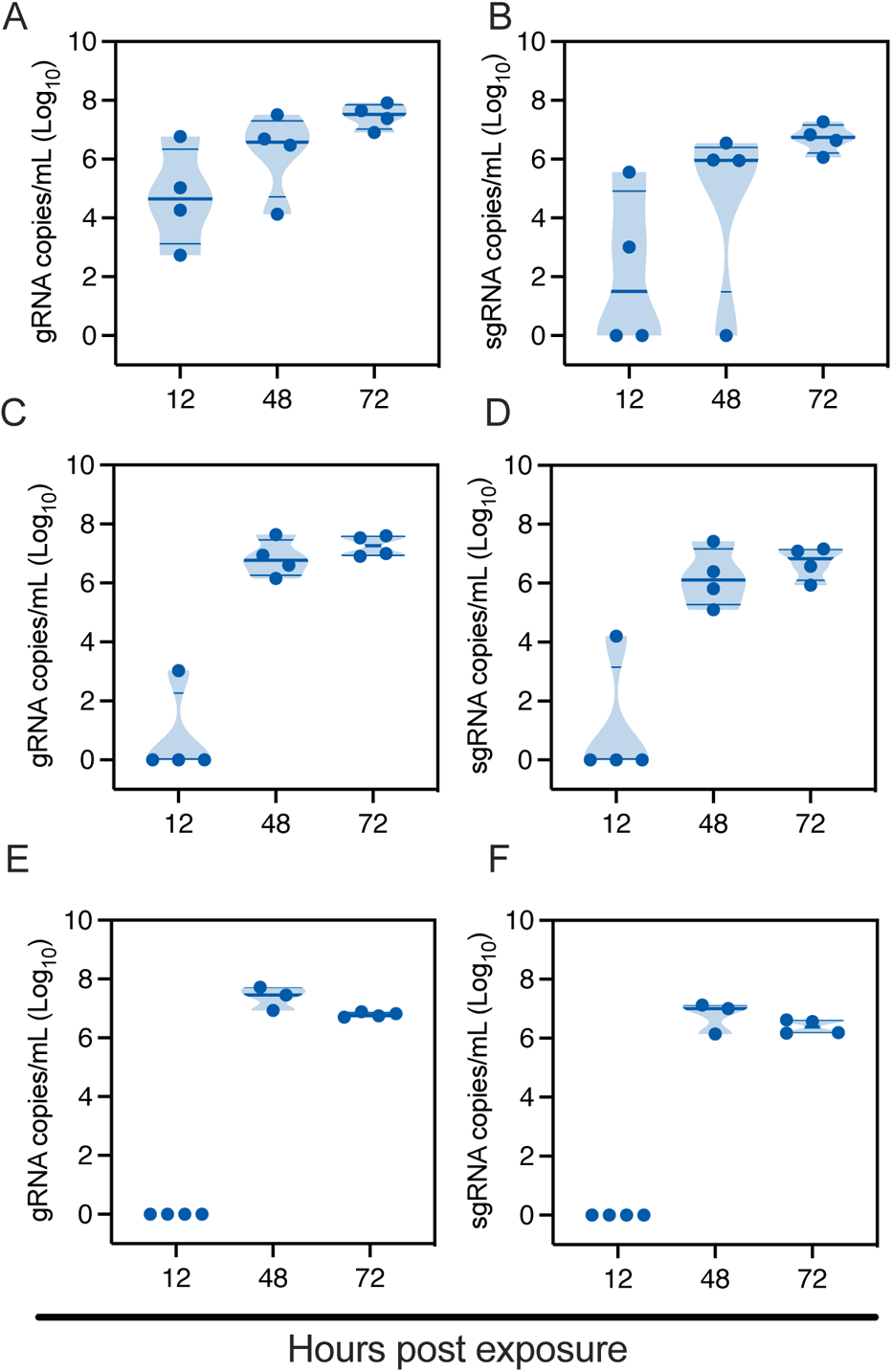
SARS-CoV-2 lineage A variant transmits efficiently over 200 cm distance. Donor Syrian hamsters were inoculated with 8×10^4^ TCID_50_ SARS-CoV-2. After 12 hours, donors were introduced to the upstream cage and sentinels (2:2 ratio) into the downstream cage. Exposure was continued for three days. To demonstrate transmission, sentinels were monitored for start and continuation of respiratory shedding. Viral load in oropharyngeal swabs of sentinels was measured by gRNA and sgRNA collected at 12, 24 and 48 hours post exposure to the donors. **A/B.** Exposure at 16.5 cm distance. **C/D.** Exposure at 106 cm distance. **E/F.** Exposure at 200 cm distance. Truncated violin plots depicting median, quantiles and individual, N = 4, two-way ANOVA, followed by Sidak’s multiple comparisons test. Abbreviations: g, gnomic; sg, subgenomic.

### Increased binding to Syrian hamster ACE2 and increased respiratory shedding of **B.1.1.7 variant**

Epidemiological studies have indicated that the emergence of the B.1.1.7 (alpha variant) was due to increased transmission over preceding virus lineages ^17^. To investigate whether the increased transmission potential of B.1.1.7 on the population level is determined by changes in transmission potential at the individual level, we compared the airborne transmission kinetics of the B.1.1.7 with the prototype lineage A virus.

First, we assessed the suitability of the Syrian hamster as a model to compare SARS-CoV-2 variant transmission. The B.1.1.7 spike binds with greater affinity to the human ACE2, potentially explaining the increased transmission ^18^. Therefore, we compared the *in-silico* binding efficiency of spike receptor binding domain (RBD) of a lineage A and B.1.1.7 variant with human and hamster ACE2. At position 501 of the B.1.1.7 spike RBD, the asparagine residue is substituted by tyrosine, which allows increased interactions with residues on ACE2 through stacking of aromatic sidechains and hydrogen bonding, hence higher affinity binding to human ACE2 ^19^. A sequence alignment between human and hamster ACE2 revealed variation at the amino acid level, however only two residues differ within the interface with SARS-CoV-2 RBD. At positions 34 and 82, histidine and methionine are replaced by glutamine and asparagine, respectively, in the hamster ACE2 (**Figure 3 A, B**). These substitutions are not located in the immediate vicinity of the N501Y mutation, suggesting that B.1.1.7 should also exhibit higher affinity binding to hamster ACE2.

**Figure 3:**
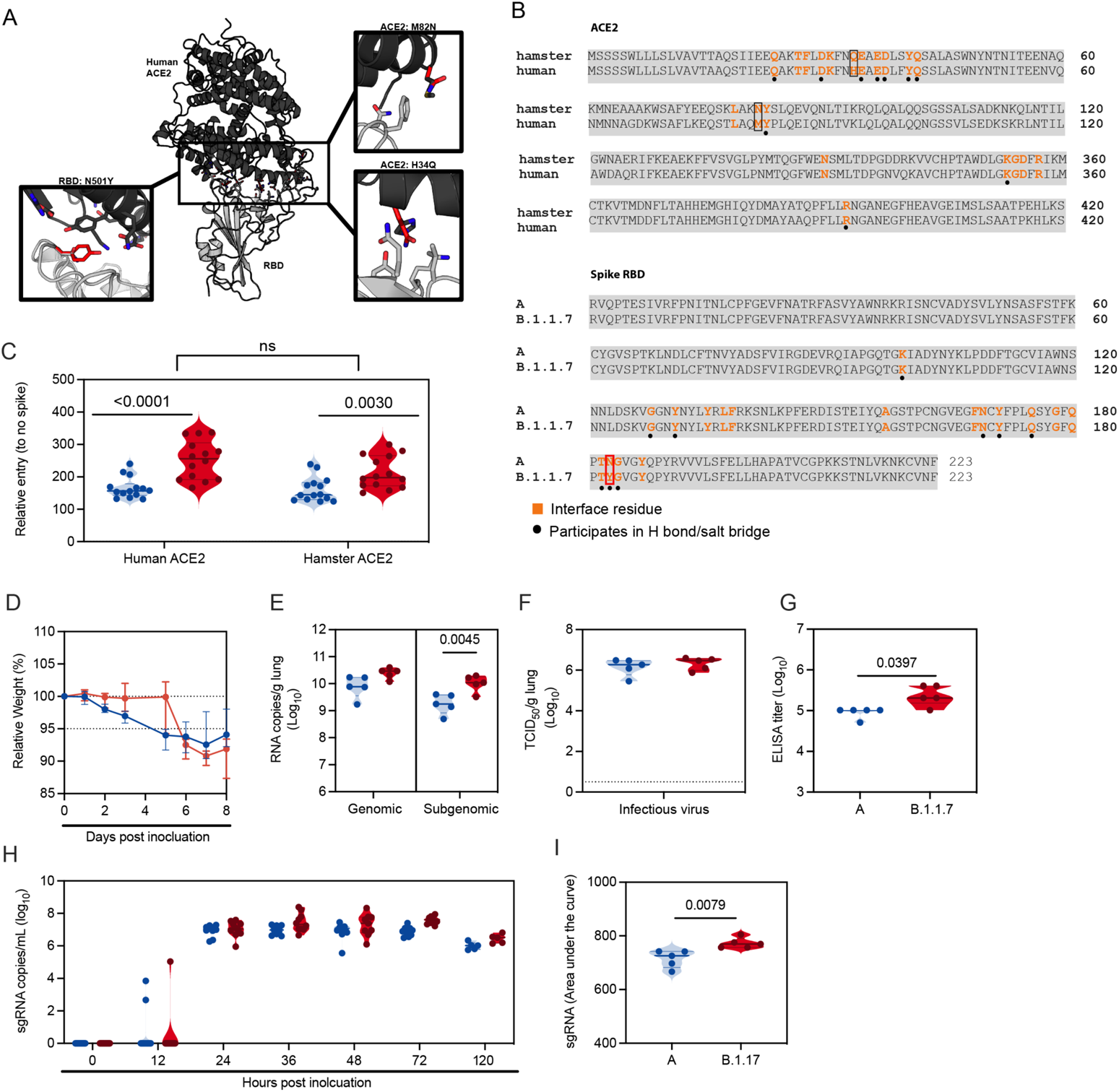
B.1.1.7 infection in Syrian hamsters is comparable to lineage A variant infection. We compared lineage A variant and B.1.1.7 receptor binding to hamster and human ACE2 *in silico* and *in vitro*. For *in vivo* comparison, Syrian hamsters were inoculated with 10^2^ TCID_50_ via the intranasal route. **A.** Differences between hamster and human ACE2 and between lineage A and B.1.1.7 SARS-CoV-2 RBD are shown on the structure of the human ACE2-RBD complex (PDB 6M0J, ^50^). The structure is shown with cartoon representation with human ACE2 colored black and RBD colored gray. Sidechains of the differing residues and surrounding residues involved in the interface, as defined by Lan *et al.* ^50^ are shown as sticks. The boxes show close up views highlighting residues that differ between the two RBDs and between human and hamster ACE2 within the interface, which were modelled using COOT ^51^. The side chains of residues at the N501Y substitution in the B.1.1.7 variant RBD, as well as the hamster ACE2 H34Q and M82N substitutions are colored red and shown superposed to the sidechain of the original residue. **B.** Amino acid sequence alignments of human ACE2 (BAB40370.1) and hamster ACE2 (XP_005074266.1), and of SARS-CoV-2 RBD from the A lineage strain and B.1.1.7 (bottom) generated using Clustal Omega. Residues involved in the RBD-ACE2 interaction, as defined by Lan, et al, ^50^, are noted in orange. Residues that participate in intermolecular hydrogen bonding or salt bridges are marked with a black dot. ACE2 residues that differ between hamster and human within the interface are outlined with a box and highlighted in (A). RBD residue 501, which differs between the A lineage variant and B.1.1.7 isolate, is also highlighted with a red box. **C**. BHK cells expressing either human ACE2 or hamster ACE2 were infected with pseudotyped VSV reporter particles with the spike proteins of WA1 and B.1.1.7 and B.1.351, luciferase was measured and normalized to no spike controls as a readout for cell entry Relative entry to no spike control for human and hamster ACE2 is depicted. Truncated violin plots depicting median, quantiles and individual, N = 14, Mann-Whitney test). **D.** Relative weight loss in hamsters after lineage A or B.1.1.7 variant inoculation. Graph shows median and 95% CI, N = 10. **E.** Viral load as measured by gRNA and sgRNA in lungs collected at day 5 post inoculation. Truncated violin plots depicting median, quantiles and individual, N = 5, ordinary two-way ANOVA followed by Sidak’s multiple comparisons test. **F.** Infectious virus determined by titration in lungs collected at day 5 post inoculation. Truncated violin plots depicting median, quantiles and individual, N = 5. **G.** Binding antibodies against spike protein of SARS-CoV-2 in serum obtained 14 days post inoculation. Truncated violin plots depicting median, quantiles and individual, N = 5, Mann-Whitney test. ELISA was performed once. **H.** Viral load as measured by sgRNA in oropharyngeal swabs collected at 0, 12, 24, 36, 48, 72 and 120 hours post inoculation. Truncated violin plots depicting median, quantiles and individual, N = 10. **I.** Area under the curve (AUC) analysis of cumulative respiratory shedding as measured by viral load in swabs. Truncated violin plots depicting median, quantiles and individual, N = 5, Mann-Whitney test, blue = lineage A, red = B.1.1.7, N. P-values are indicated were appropriate. Abbreviations: A, lineage A variant; g, gnomic; sg, subgenomic; RBD, receptor binding domain; ACE2, Angiotensin-converting enzyme 2.

To confirm that the observed enhanced binding affinity of B.1.1.7 to human ACE2 was also present for hamster ACE2 we directly compared viral entry using a VSV pseudotype entry assay. No significant difference in entry between human and hamster ACE2 with either lineage A or B.1.1.7 was observed. For both human and hamster ACE2, B.1.1.7 demonstrated significantly increased entry compared to the lineage A variant (human ACE2 median lineage A/B.1.1.7 = 156.8/256 (relative entry to no spike), p <0.0001 and hamster ACE2 median lineage A/B.1.1.7 = 144.6/197.5 (relative entry to no spike), p = 0.003, N = 14, Mann-Whitney test) (**Figure 3 C**).

We next investigated if the enhanced binding affinity of B.1.1.7 to hamster ACE2 translated to differences in viral replication and shedding dynamics *in-vivo*. Hamsters (N = 10) were inoculated I.N. with 10^2^ TCID_50_ of SARS-CoV-2 lineage A variant (A) or B.1.1.7. Regardless of variant, weight loss was observed in all animals with a maximum at 7 days post inoculation (DPI), after which animals began to recover (**Figure 3 D**, N = 5, median weight loss lineage A/B.1.1.7 = 7.5/8.2%). Five out of ten hamsters per group were euthanized at 5 DPI and lung tissue was harvested to assess viral replication in the lower respiratory tract. Lung tissue of animals inoculated with B.1.1.7 contained higher levels of gRNA and significantly higher levels of sgRNA (**Figure 3 E**, N = 5, ordinary two-way ANOVA, followed by Sidak’s multiple comparison test, median lineage A/B.1.1.7 = 9.9/10.5 log_10_ copies/g and p = 0.0614; median lineage A/B.1.1.7 = 9.2/10 log_10_ copies/g and p = 0.0045, respectively). Infectious virus titers in the lungs were not significantly different between the variants (**Figure 3 F**, N = 5 median lineage A/B.1.1.7 = 6.3/6.5 log_10_ TCID_50_/g). At 14 DPI, all remaining animals had seroconverted. Anti-spike IgG ELISA titers were significantly increased in animals inoculated with B.1.1.7 (**Figure 3 G**, N = 5, Mann-Whitney test, median lineage A/B.1.1.7 = 102400/204800 and p = 0.0394). Next, we studied differences in shedding from the upper respiratory tract. sgRNA could be detected in two animals at 12 hours after inoculation (HPI) with lineage A variant and in one animal inoculated with B.1.1.7. sgRNA was detected at similar levels for both groups at 24 HPI (**Figure 3 H**). At 3 DPI, a significant increase in sgRNA was seen in B.1.1.7 inoculated animals. In both groups sgRNA levels from oropharyngeal swabs started to drop at 5 DPI, levels in the B.1.1.7 animals remained somewhat higher (3 DPI median lineage A/B.1.1.7 = 6.9/7.6 and 5 DPI median lineage A/B.1.1.7 = 6.0/6.5 copies/mL (log_10_)). This translated to a significant difference when comparing the cumulative shedding until 5 DPI (**Figure 3 I**, area under the curve (AUC), N = 5, Mann-Whitney test, median lineage A/B.1.1.7 = 726/770 cumulative copies/mL (log_10_) and p = 0.0079).

### Efficient aerosol transmission with B.1.1.7

We repeated the aerosol transmission experiment at 106 cm and 200 cm as described above for B.1.1.7. Aerosol transmission of B.1.1.7 was equally as efficient as for lineage A; all sentinels demonstrated respiratory shedding and seroconversion (**Figure 4 A/B/C/D, Supplementary Table 1**).

**Figure 4:**
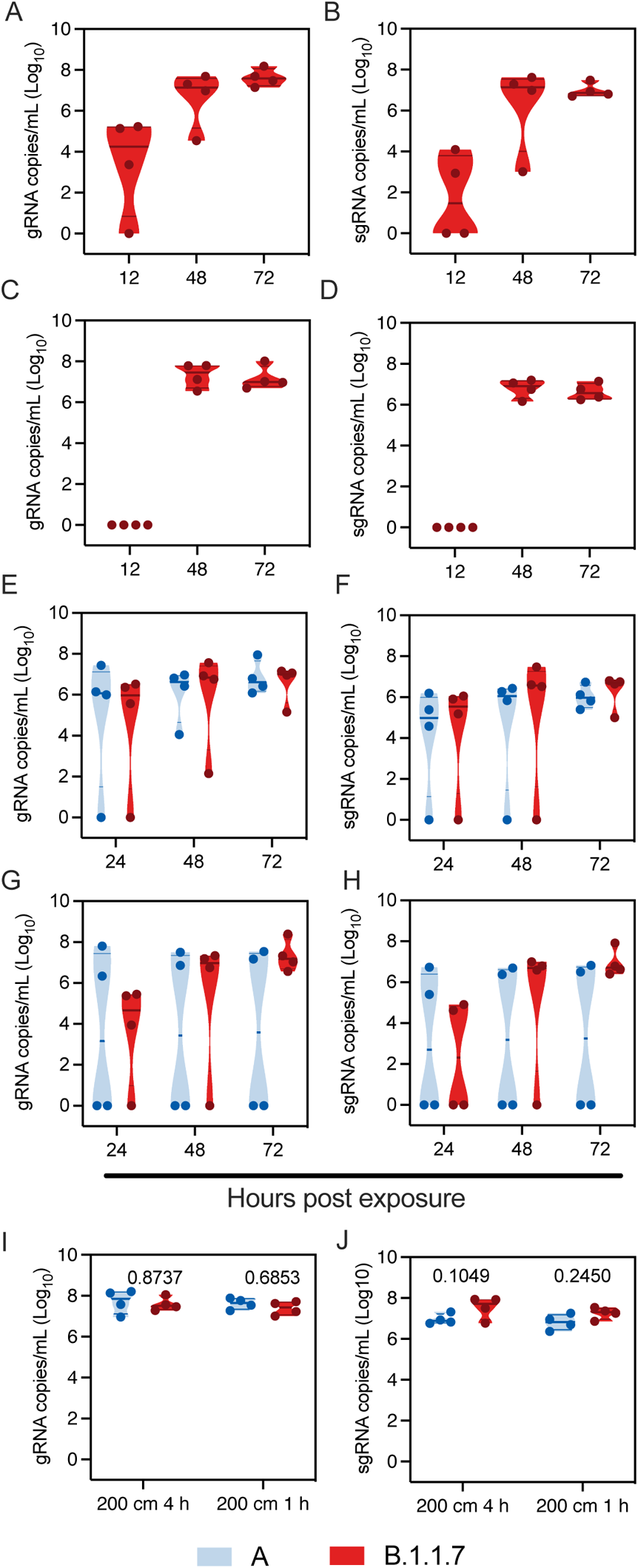
B.1.1.7 aerosol transmission efficiency is increased. Comparison of aerosol transmission efficiency of lineage A and B.1.1.7 SARS-CoV-2 variants in the Syrian hamster. **A/B.** Donor Syrian hamsters were inoculated with 8×10^4^ TCID_50_ SARS-CoV-2 B.1.1.7 variant. After 12 hours, donors were introduced to the upstream cage and sentinels (2:2 ratio) into the downstream cage. Exposure was continued for three days. To demonstrate transmission, sentinels were monitored for start and continuation of respiratory shedding. Viral load in oropharyngeal swabs of sentinels was measured by gRNA and sgRNA collected at 12, 24 and 48 hours post exposure to the donors. Exposure at 106 cm distance. **C/D.** Exposure at 200 cm distance. **E/F.** Donor Syrian hamsters were inoculated with 8×10^4^ TCID_50_ SARS-CoV-2 B.1.1.7 variant or lineage A (N = 4, respectively). After 12 hours, donors were introduced to the upstream cage and sentinels (2:2 ratio) into the downstream cage. Exposure was limited to four hours. gRNA and sgRNA in oropharyngeal swabs were collected at 24, 48 and 72 hours post exposure to the donors. **G/H.** Exposure was limited to one hour for B.1.1.7 and lineage A (N = 4, respectively). gRNA and sgRNA in oropharyngeal swabs were collected at 24, 48 and 72 hours post exposure to the donors. **I/J.** Viral load in oropharyngeal swabs of donors was measured by gRNA and sgRNA collected 24 hours post inoculation. Truncated violin plots depicting median, quantiles and individual, blue = lineage A, red = B.1.1.7, N = 4 for each variant, two-way ANOVA, followed by Sidak’s multiple comparisons test. P-values are indicated were appropriate. Abbreviations: A, lineage A variant; g, gnomic; sg, subgenomic.

We next set out to determine the transmission efficiency of both lineage A and B.1.1.7 within a limited exposure window of either one or four hours at 200 cm distance. First, four donor animals were I.N. inoculated with 8×10^4^ TCID_50_ SARS-CoV-2 lineage A and four donors with B.1.1.7. Sentinel were exposed in a 2:2 ratio at 12 hours post inoculation for a duration of four hours. Transmission via aerosols occurred even when time of exposure was limited for both variants. Both gRNA and sgRNA shedding were detected 24 hours after exposure in oropharyngeal swabs of three out of four sentinel animals exposed to B.1.1.7 and lineage A (**Figure 4 E/F**). At 3 DPE, all sentinels displayed gRNA and sgRNA shedding.

Next, the same experiment was repeated with the exposure time limited to one hour. gRNA was detected 24 hours post exposure in oropharyngeal swabs of three out of four sentinels exposed to B.1.1.7 and two out of four sentinels exposed to lineage A, respectively (**Figure 4 G/H**). sgRNA was detected in oropharyngeal swabs of two out of four sentinels exposed to B.1.1.7 and two out of four sentinels for lineage A, respectively. At 3 DPE, whilst all sentinels exposed to B.1.1.7 were positive for gRNA and sgRNA, viral RNA was only detected in two of four sentinels exposed to lineage A. Viral loads in swabs did not differ significantly between the two variants. To ensure the differences observed in transmission were not due to increased donor shedding, we compared viral loads in oropharyngeal swabs taken from donor animals after exposure. B.1.1.7 did not impact the respiratory shedding of the donors at this time point (**Figure 4 I/J**, N = 4, ordinary two-way ANOVA, followed by Sidak’s multiple comparison test, four hours: gRNA p = 0.8737, sgRNA p = 0.1049, one hour: gRNA p = 0.6853, sgRNA p = 0.2450). The same experiment was repeated on day three after inoculation of the donors. No transmission occurred at this time point; we did not observe gRNA in oropharyngeal swabs of any sentinel on consecutive days and no sgRNA was detected in any swab taken during three days post exposure (**Supplementary Figure 2**). These data suggest that aerosol transmission for B.1.1.7 may be more efficient compared to lineage A, and is independent of amount of virus shed by donor.

### B.1.1.7 variant demonstrates increased airborne transmission competitiveness

To assess the transmission efficiency in direct competition between lineage A and B.1.1.7 variants, we employed the 16.5 cm cage system to conduct a transmission chain study. Donor animals (N = 8) were inoculated I.N. with 1×10^2^ TCID_50_ SARS-CoV-2 (1:1 lineage A:B.1.1.7 mixture). Dual infection presented with comparable weight loss and shedding profile to inoculation with either variant (**Supplementary Figure 3 A/B**). After 12 hours donors were co-housed with eight sentinels (Sentinels 1) (2:2 ratio) for 24 hours (**Figure 5 A**). Immediately after, the eight sentinels were co-housed with eight new sentinels (Sentinels 2) (2:2 ratio) for 24 hours and donor animals were relocated to normal rodent caging. This sequence was repeated for Sentinels 3. For each round, the previous sentinels were housed in the upstream cage and became the new donors. We assessed transmission by measuring viral RNA in oropharyngeal swabs taken from all animals at 2 DPI/DPE. While all donor animals (median gRNA = 7.3 copies/mL (log_10_), median sgRNA = 7.0 copies/mL (log_10_)) and all Sentinels 1 (median gRNA = 7.0 copies/mL (log_10_), median sgRNA = 6.8 copies/mL (log_10_)) demonstrated robust shedding, viral RNA could only be detected in four out of eight Sentinels 2 (median gRNA = 2.5 copies/mL (log_10_), median sgRNA = 1.8 copies/mL (log_10_)), and in one Sentinels 3 animal (**Figure 5 B/C**). We compared infectious virus titers in the swabs. While all donor animals (median = 4.25 TCID_50_ (log_10_) and all Sentinels 1 had high infectious virus titers (median = 4.5 TCID_50_ (log_10_)), infectious virus could only be detected in four Sentinels 2 (median = 0.9 TCID_50_ (log_10_)), and no Sentinels 3 animals (**Figure 5 D**). We then proceeded to compare the viral loads in the lungs of these animals at 5 DPE. As expected, viral RNA was only detected in animals that were positive for SARS-CoV-2 in their corresponding oropharyngeal swab. While all donor animals (median sgRNA = 10.0 copies/g (log_10_)) and all Sentinels 1 had high gRNA levels in the lung (median sgRNA = 10.0 copies/mL (log_10_), viral RNA could only be detected in four Sentinels 2 (median sgRNA = 4.3 copies/mL (log_10_), and no RNA was detected in any Sentinels 3 (**Figure 5 E**). We compared the gross pathology of these lungs. Lungs from Sentinels 1 demonstrated SARS-CoV-2 infection associated pathology as previously described ^14,20,21^. Pathology was only seen in three Sentinels 2 and no Sentinels 3 (**Figure 4 F, Supplementary Table 3, Supplementary Figure 3 C**). This suggests that transmission very early after exposure may be restricted and that not all animals were able to efficiently transmit the virus to the next round of naïve sentinels.

**Figure 5:**
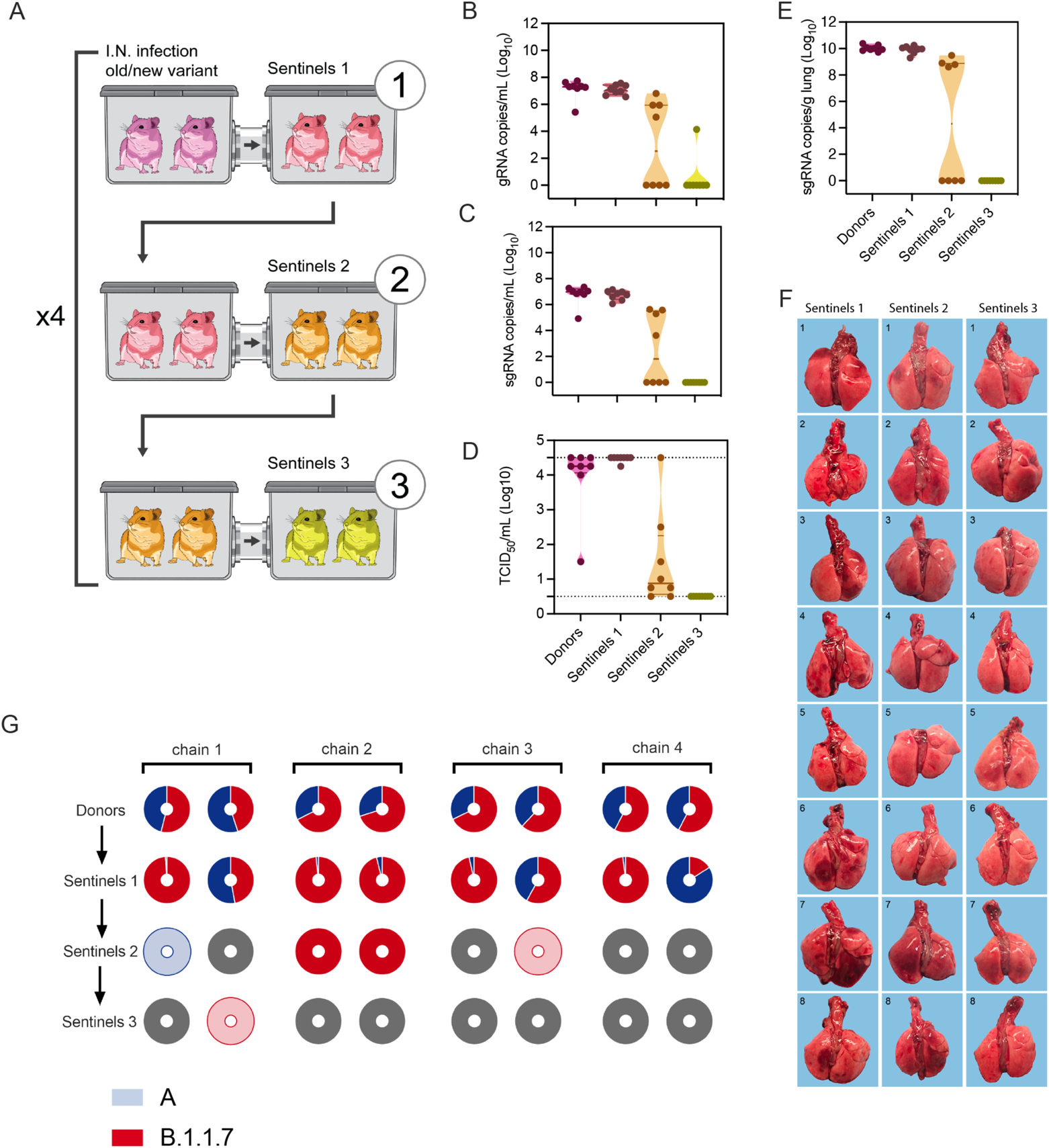
B.1.1.7 variant has increased airborne transmission competitiveness. Donor animals (N = 8) were inoculated with both lineage A and B.1.1.7 variant with 10^2^ TCID_50_ via the intranasal route (1:1 ratio), and three groups of sentinels (Sentinels 1, 2 and 3) were exposed subsequently at 16.5 cm distance. **A**. Schematic visualization of the transmission chain design. Animals were exposed at a 2:2 ratio, exposure occurred on consecutive days and lasted for 24 hours for each chain link. **B/C.** Respiratory shedding measured by viral load in oropharyngeal swabs; measured by gRNA and sgRNA on day 2 post exposure. Truncated violin plots depicting median, quantiles and individuals, N = 8. **D.** Corresponding infectious virus in oropharyngeal swabs, measured by titration. Truncated violin plots depicting median, quantiles and individuals, N = 8. **E.** Corresponding infectious virus in lungs sampled five days post exposure, measured by titration. Truncated violin plots depicting median, quantiles and individuals, N = 8. **F.** Gross pathology of lungs at day 5 post exposure. **G.** Percentage of B.1.1.7 detected in oropharyngeal swabs taken at day 2 post exposure for each individual donor and sentinel, determined by deep sequencing. Pie-charts depict individual animals. Red = B.1.1.7, blue = lineage A, grey = no viral RNA present in sample, and transparent color = duplex-qRT-PCR confirmed viral RNA presence in sample but sequencing unsuccessful due to low RNA quality.

To determine the competitiveness of the variants we analyzed the relative composition of the two viruses in the swabs using next generation sequencing and compared the percentage of B.1.1.7/lineage A at 2 DPE (**Figure 5 G, Supplementary Table 4**). We observed one donor with increased amounts of lineage A variant (55%), while in the remaining seven animals the B.1.1.7 variant was increased (54-74% range). After the first airborne transmission sequence, two sentinels shed increased amounts of lineage A variant (55% and 84%), while the remaining six shed more B.1.1.7; five of which shed nearly exclusively B.1.1.7 (>96%). After the second round of the airborne transmission sequence, three out of four sentinel animals shed exclusively B.1.1.7 and one animal shed exclusively lineage A. Due to low amounts of viral RNA, two sentinels in the Sentinels 2 and the one sentinel in the Sentinels 3 group could not be successfully sequenced. We analyzed these samples by duplex-qRT-PCR applying a modified 2^−ΔΔCt^ method (**Supplementary Table 4**). One animal in the Sentinels 2 group only shed B.1.1.7 (no lineage A PCR positivity). The other shed nearly exclusively lineage A (0.0007-fold increase of B.1.1.7), while, interestingly, the transmission event of this animal to the Sentinels 3 animal was exclusively B.1.1.7. Taken together, B.1.1.7 demonstrated increased competitiveness; in 10 out of 13 airborne transmission events B.1.1.7 outcompeted lineage A and only in three events infection with lineage A was established as the dominant variant.

## Discussion

Epidemiological studies in humans strongly suggest that aerosol transmission plays a major role in driving the SARS-CoV-2 pandemic ^22–24^. Yet formal proof of aerosol transmission of SARS-CoV-2 has not been provided and would rely on demonstration of long distance transmission in the absence of other transmission routes ^25^. Here we demonstrated efficient transmission of SARS-CoV-2 between Syrian hamsters via particles <5 µm ^6^ over 200 cm distance. Additionally, we present first qualitative analyses of the efficiency of transmission, showing that even within one hour transmission can occur at a distance of 200 cm between Syrian hamsters. Whereas several SARS-CoV-2 transmission studies in hamsters and ferrets have been performed, none of these studies were able to differentiate between large and small droplet transmission ^14,15,26–28^. In our previous work, specifically designed cage dividers were used to generate an airflow system minimizing large droplet cross-over. While the number of large droplets was markedly reduced, aerosol transmission could not be conclusively demonstrated ^14^. Within the currently described transmission caging only 2% and 0.5% of particles found in the sentinel side were ≥5 µm at 106 and 200 cm distance, respectively, strongly suggesting that the transmission observed in these cages is by true aerosols. This is an important finding in two regards. First, epidemiological conclusive evidence for aerosol transmission of SARS-CoV-2 is currently still lacking, because it is difficult to determine with certainty the route or combination of routes of transmission. Second, particles <5 µm are expected to reach the respiratory bronchioles and alveoli. While respirable aerosol (<2.5 µm), thoracic aerosol (<10 µm) and inhalable aerosol in general ^29^ all may be relevant to infection with SARS-CoV-2 ^30^, it has been suggested that direct deposition into the lower respiratory tract may decrease the necessary infectious dose ^31^. Indeed, our previous work has demonstrated that aerosol inoculation in the Syrian hamster is highly efficient (25 TCID_50_, particles <5µm ^32^) and is linked to increased disease severity due to direct deposition of the virus into the lower respiratory tract ^14^.

The data presented here need to be considered in the context of inherent differences between the Syrian hamster model and human behavior. Experimentally, animals were exposed to a unidirectional airflow at timepoints chosen for optimal donor shedding, which likely contributed to the high efficiency of aerosol transmission even at 200 cm distance after only one hour of exposure. However, this approximates human exposure settings such as restaurants or office spaces.

Increased risk of airborne transmission is an important concern in the context of VOCs. VOC B.1.1.7 was first detected in the United Kingdom and has been shown to exhibit increased transmission with significantly increased reproduction number and attack rates ^33,34^. Transmission efficiency is a function of donor shedding, exposure time, sentinel susceptibility, and potential environmental factors effecting stability during transmission. One experimental study in preprint has found no difference in contact, fomite or short distance airborne transmission between D614G and B.1.1.7 in the Syrian hamster, however, numbers were low and no aerosol transmission was compared ^35^. Our entry data shows that there is increased entry of B.1.1.7 over lineage A for both human and hamster ACE2, confirming the suitability of the Syrian hamster model for variant comparison. Our data suggest that the increased transmission efficiency of B.1.1.7 may not be a direct result of the shedding magnitude but that a lower dose of B.1.1.7 may be sufficient for transmission. Under the applied experimental restrictions (200 cm, 1 hour) both lineage A and B.1.1.7 transmitted equally as efficient. However, B.1.1.7 displayed an increased airborne transmission competitiveness over lineage A in a dual infection experiment. This has also previously been shown for D614G over the lineage A variant ^36,37^ and for B.1.1.7 over D614G ^38^, however, these studies did not look at airborne transmission. Previously, the D614G competition and transmission experiments in hamsters and ferrets suggested that the D614G mutation increased transmissability ^39,40^. The additional N501Y mutation is specifically predicted to increase affinity for human ACE2, partially explaining the dominance of B.1.1.7 and other new variants containing both mutations ^41,42^. Additional work is required to demonstrate conclusively if the increased airborne competitiveness of B.1.1.7 in the hamster model is truly a result of increased susceptibility due to better viral entry and/or decreased infectious dose. Our data indicate that the Syrian hamster represents a valuable model to rapidly evaluate transmission differences between novel VOCs

The increase in aerosol transmission potential of B.1.1.7 underscores the continuous need for development and implementation of non-pharmaceutical preemptive interventions. In the light of limited global vaccine coverage and the potential emergence of escape mutants, ventilation, and air disinfection ^43,44^, face masks and social distancing ^13,45^, should still be considered essential tools in COVID-19 exposure and transmission risk mitigation strategies.

## Materials and Methods

### Ethics Statement

All animal experiments were conducted in an AAALAC International-accredited facility and were approved by the Rocky Mountain Laboratories Institutional Care and Use Committee following the guidelines put forth in the Guide for the Care and Use of Laboratory Animals 8th edition, the Animal Welfare Act, United States Department of Agriculture and the United States Public Health Service Policy on the Humane Care and Use of Laboratory Animals. Work with infectious SARS-CoV-2 virus strains under BSL3 conditions was approved by the Institutional Biosafety Committee (IBC). For the removal of specimens from high containment areas virus inactivation of all samples was performed according to IBC-approved standard operating procedures.

### Cells and virus

SARS-CoV-2 variant B.1.1.7 (hCoV320 19/England/204820464/2020, EPI_ISL_683466) was obtained from Public Health England via BEI. SARS-CoV-2 strain nCoV-WA1-2020 (lineage A, MN985325.1) was provided by CDC, Atlanta, USA. Virus propagation was performed in VeroE6 cells in DMEM supplemented with 2% fetal bovine serum, 1 mM L-glutamine, 50 U/mL penicillin and 50 μg/mL streptomycin (DMEM2). VeroE6 cells were maintained in DMEM supplemented with 10% fetal bovine serum, 1 mM L-glutamine, 50 U/mL penicillin and 50 μg/ml streptomycin. At regular intervals mycoplasma testing was performed. No mycoplasma and no contaminants were detected. For sequencing from viral stocks, sequencing libraries were prepared using Stranded Total RNA Prep Ligation with Ribo-Zero Plus kit per manufacturer’s protocol (Illumina) and sequenced on an Illumina MiSeq at 2 × 150 base pair reads. No nucleotide change was found >5% for nCoV-WA1-2020, while for VOC B.1.1.7 **Supplementary Table 6** summarized the mutations obtained.

### Plasmids

The spike coding sequences for SARS-CoV-2 lineage A (nCoV-WA1-2020) and variant B.1.1.7 (hCoV320 19/England/204820464/2020, EPI_ISL_683466) were truncated by deleting 19 aa at the C-terminus. The S proteins with the 19 aa deletion of coronaviruses were previously reported to show increased efficiency regarding incorporation into virions of VSV ^46,47^. These sequences were codon optimized for human cells, then appended with a 5′ kozak expression sequence (GCCACC) and 3′ tetra-glycine linker followed by nucleotides encoding a FLAG-tag sequence (DYKDDDDK). These spike sequences were synthesized and cloned into pcDNA3.1^+^(GenScript). Human and hamster ACE2 (Q9BYF1.2 and GQ262794.1, respectively), were synthesized and cloned into pcDNA3.1^+^ (GenScript). All DNA constructs were verified by Sanger sequencing (ACGT).

### Receptor transfection

BHK cells were seeded in black 96-well plates and transfected the next day with 100 ng plasmid DNA encoding human or hamster ACE2, using polyethylenimine (Polysciences). All downstream experiments were performed 24 h post-transfection.

### Pseudotype production and Luciferase-based cell entry assay

Pseudotype production was carried as described previously ^48^. Briefly, plates pre-coated with poly-L-lysine (Sigma–Aldrich) were seed with 293T cells and transfected the following day with 1,200 ng of empty plasmid and 400 ng of plasmid encoding coronavirus spike or no-spike plasmid control (green fluorescent protein (GFP)). After 24 h, transfected cells were infected with VSVΔG seed particles pseudotyped with VSV-G, as previously described ^48,49^. After one hour of incubating with intermittent shaking at 37 °C, cells were washed four times and incubated in 2 mL DMEM supplemented with 2% FBS, penicillin/streptomycin and L-glutamine for 48 h. Supernatants were collected, centrifuged at 500x*g* for 5 min, aliquoted and stored at −80 °C. BHK cells previously transfected with ACE2 plasmid of interest were inoculated with equivalent volumes of pseudotype stocks. Plates were then centrifuged at 1200x*g* at 4 °C for one hour and incubated overnight at 37 °C. Approximately 18–20 h post-infection, Bright-Glo luciferase reagent (Promega) was added to each well, 1:1, and luciferase was measured. Relative entry was calculated normalizing the relative light unit for spike pseudotypes to the plate relative light unit average for the no-spike control. Each figure shows the data for two technical replicates.

### Structural interaction analysis

Structure modeling was performed using the human ACE2 and SARS-CoV-2 RBD crystal structure, PDB ID 6M0J ^50^. Mutagenesis to model the residues that differ in the B.1.1.7 RBD and hamster ACE2 was performed in COOT ^51^. The structure figure was generated using the Pymol Molecular Graphics System (https://www.schrodinger.com/pymol). Amino acid sequence alignments of human ACE2 (BAB40370.1) and hamster ACE2 (XP_005074266.1), and of SARS-CoV-2 RBD from the linage A strain and B.1.1.7 variant, were were generated using Clustal Omega (http://europepmc.org/article/MED/). Residues participating in the SARS-CoV-2 – ACE2 interface were noted as described by Lan, et al ^50^.

### Duplex-qRT-PCR variant detection

Duplex-qRT-PCR primers and probe were designed to distinguish between lineage A SARS-CoV-2 and B.1.1.7 variant (**Supplemental Table 5**) in a duplex assay. The forward and reverse primers were design to detect both variants while two probes were designed to detect either variant. Five μL RNA was tested with TaqMan™ Fast Virus One-Step Master Mix (Applied Biosystems) using QuantStudio 3 Real-Time PCR System (Applied Biosystems) according to instructions of the manufacturer. Relative fold-change difference between both variants was calculated by applying the delta-delta Ct method, (2^−ΔΔCt^ method) with modifications.

### Inoculation experiments

Four to six-week-old female and male Syrian hamsters (ENVIGO) were inoculated (10 animals per virus) intranasally (I.N) with either SARS-CoV-2 strain nCoV-WA1-2020 (lineage A) or hCoV320 19/England/204820464/2020 (B.1.1.7), or a 1:1 mixture of both viruses. I.N. inoculation was performed with 40 µL sterile DMEM containing 1×10^2^ TCID_50_ SARS-CoV-2. At five days post inoculation (DPI), five hamsters for each route were euthanized, and tissues were collected. The remaining 5 animals for each route were euthanized at 14 DPI for disease course assessment and shedding analysis. Hamsters were weighted daily, and oropharyngeal swabs were taken on day 1, 2, 3 and 5. Swabs were collected in 1 mL DMEM with 200 U/mL penicillin and 200 µg/mL streptomycin. Hamsters were observed daily for clinical signs of disease. Necropsies and tissue sampling were performed according to IBC-approved protocols.

### Aerosol cages

The aerosol transmission system consisted of two 7” X 11” X 9” plastic hamster boxes (Lab Products, Inc.) connected with a 3” diameter tube (**Supplementary Figure 1**). The boxes were modified to accept a 3” plastic sanitary fitting (McMaster-Carr) which enabled the length between the boxes to be changed. The nominal tube lengths were 16.5, 106 and 200 cm. Airflow was generated with a vacuum pump (Vacuubrand) attached to the box housing the naïve animals and was controlled with a float type meter/valve (King Industries, McMaster-Carr). The airflow was adjusted for each tube length to be 30 cage changes/hour and the flow was validated prior to starting the experiments by timing a smoke plume through the tubes. The airflow of the original boxes is in through a filtered top and out through an exhaust port in the side of the box. To ensure proper airflow from the donor box to the naïve box, the top of the naïve box was sealed while the filter top of the donor box remained open.

To ensure the system was able to contain aerosols the airtightness of the system was validated with a negative pressure smoke test and a positive pressure leak test prior to moving into a containment laboratory. To perform the negative pressure test the airflow was adjusted to exhaust the system at 30 cage changes/hour, smoke was generated in the donor cage with a WizardStick and escaped particulate was measured with a TSI DustTrak DRX. To test the system under pressure the air flow was reversed, and the joints were tested using a gas leak detector.

### Particle sizing

Transmission cages were modified by introducing an inlet on the side wall of the infected hamster side, and sample ports on each end of the connection tube for measurement of particles in the air under constant airflow condition. Particles were generated by spraying a 20% (v/v) glycerol solution with a standard spray bottle through the donor cage inlet. The particle size was measured using a Model 3321 aerodynamic particle sizer spectrometer (TSI). First, the donor cage was coated with three sprays at an interval of 30 seconds (s). The sample port was opened, and a sample was analyzed. Every 30 s a new spray followed, and five samples were analyzed (5 runs, each 60 s) for both donor side (primary infected side) and sentinel side.

### Aerosol Transmission experiments

All transmission studies were conducted at a 2:2 ratio between donor and sentinels for each transmission scenario tested and virus variant with 2 separate transmission cages (N = 4 donors / 4 sentinels). To ensure no cross-contamination, the donor cages and the sentinel cages were never opened at the same time, sentinel hamsters were not exposed to the same handling equipment as donors and after each sentinel the equipment was disinfected with either 70% ETOH or 5% Microchem.

Initially, transmission was studied assessing distance. Donor hamsters were infected intranasally as described above with 8×10^4^ TCID_50_ SARS-CoV-2 (lineage A or B.1.1.7 variants). After 12 hours donor animals were placed into the donor cage and sentinels were placed into the sentinel cage (2:2). Air flow was generated between the cages from the donor to the sentinel cage at 30 changes/h. Hamsters were co-housed at 16.5 cm, 106 cm or 200 cm distance. Regular bedding was replaced by alpha-dri bedding to avoid the generation of dust particles. Oropharyngeal swabs were taken for donors at 1 DPI and for sentinels daily after exposure began. Swabs were collected in 1 mL DMEM with 200 U/mL penicillin and 200 µg/mL streptomycin. Exposure continued until respiratory shedding was confirmed in sentinels on three consecutive days. Then donors were euthanized, and sentinels were monitored until 14 DPE (days post exposure) for seroconversion.

Second, transmission was studied assessing duration of exposure. Donor hamsters were infected intranasally as described above with 8×10^4^ TCID_50_ SARS-CoV-2. After 24 hours (1 DPI) or 72 hours (3 DPI) donor animals were placed into the donor cage and sentinels were placed into the sentinel cage (2:2). Hamsters were co-housed at 200 cm distance for 1 or 4 hours at an airflow rate of 30 changes/h. Oropharyngeal swabs were taken for donors at day of exposure and for sentinels for three days after exposure.

### Variant competitiveness transmission chain

Donor hamsters (N = 8) were infected intranasally as described above with 1×10^2^ TCID_50_ SARS-CoV-2 at a 1:1 ratio of lineage A and B.1.1.7 mixture. After 12 hours donor animals were placed into the donor cage and sentinels (Sentinels 1, N = 8) were placed into the sentinel cage (2:2) at 16.5 cm distance at an airflow of 30 changes/h. Hamsters were co-housed for 24 h. The following day, donor animals were re-housed into regular rodent caging and Sentinels 1 were placed into the donor cage of new transmission set-ups. New sentinels (Sentinels 2, N = 8) were placed into the sentinel cage (2:2) at 16.5 cm distance at an airflow of 30 changes/h. Hamsters were co-housed for 24 h. Then, Sentinels 1 were re-housed into regular rodent caging and Sentinels 2 were placed into the donor cage of new transmission set-ups. New sentinels (Sentinels 3, N = 8) were placed into the sentinel cage (2:2) at 16.5 cm distance at an airflow of 30 changes/h. Hamsters were co-housed for 24 h. Then, both Sentinels 2 and Sentinels 3 were re-housed to regular rodent caging and monitored until 5 DPE. Oropharyngeal swabs were taken for all animals at 2 DPI/DPE. All animals were euthanized at 5 DPI/DPE for collection of lung tissue.

### Viral RNA detection

Swabs from hamsters were collected as described above. Then, 140 µL was utilized for RNA extraction using the QIAamp Viral RNA Kit (Qiagen) using QIAcube HT automated system (Qiagen) according to the manufacturer’s instructions with an elution volume of 150 µL. For tissues, RNA was isolated using the RNeasy Mini kit (Qiagen) according to the manufacturer’s instructions and eluted in 60 µL. Sub-genomic (sg) viral RNA and genomic (g) was detected by qRT-PCR ^52,53^. RNA was tested with TaqMan™ Fast Virus One-Step Master Mix (Applied Biosystems) using QuantStudio 6 or 3 Flex Real-Time PCR System (Applied Biosystems). SARS-CoV-2 standards with known copy numbers were used to construct a standard curve and calculate copy numbers/mL or copy numbers/g.

### Viral titration

Viable virus in tissue samples was determined as previously described ^54^. In brief, lung tissue samples were weighted, then homogenized in 1 mL of DMEM (2% FBS). VeroE6 cells were inoculated with ten-fold serial dilutions of homogenate, incubated 1 hours at 37°C and the first two dilutions washed twice with 2% DMEM. After 6 days cells were scored for cytopathic effect. TCID_50_/mL was calculated by the method of Spearman-Karber.

### Serology

Serum samples were analyzed as previously described ^55^. In brief, maxisorp plates (Nunc) were coated with 50 ng spike protein (generated in-house) per well. Plates were incubated overnight at 4°C. Plates were blocked with casein in phosphate buffered saline (PBS) (ThermoFisher) for 1 hours at room temperature (RT). Serum was diluted 2-fold in blocking buffer and samples (duplicate) were incubated for 1 hours at RT. Secondary goat anti-hamster IgG Fc (horseradish peroxidase (HRP)-conjugated, Abcam) spike-specific antibodies were used for detection and visualized with KPL TMB 2-component peroxidase substrate kit (SeraCare, 5120-0047). The reaction was stopped with KPL stop solution (Seracare) and plates were read at 450 nm. The threshold for positivity was calculated as the average plus 3 x the standard deviation of negative control hamster sera.

### Next-generation sequencing of virus

For sequencing from swabs, total RNA was depleted of ribosomal RNA using the Ribo-Zero Gold rRNA Removal kit (Illumina). Sequencing libraries were constructed using the KAPA RNA HyperPrep kit following manufacturer’s protocol (Roche Sequencing Solutions). To enrich for SARS-CoV-2 sequence, libraries were hybridized to myBaits Expert Virus biotinylated oligonucleotide baits following the manufacturer’s manual, version 4.01 (Arbor Biosciences, Ann Arbor, MI). Enriched libraries were sequenced on the Illumina MiSeq instrument as paired-end 2 X 150 base pair reads. Raw fastq reads were trimmed of Illumina adapter sequences using cutadapt version 1.1227 and then trimmed and filtered for quality using the FASTX-Toolkit (Hannon Lab, CSHL). Remaining reads were mapped to the SARS-CoV-2 2019-nCoV/USA-WA1/2020 (MN985325.1 using Bowtie2 version 2.2.928 with parameters --local --no-mixed-X 1500. PCR duplicates were removed using picard MarkDuplicates (Broad Institute) and variants were called using GATK HaplotypeCaller version 4.1.2.029 with parameter-ploidy 2. Variants were filtered for QUAL > 500 and DP > 20 using bcftools. We assessed the presence of N501Y, D614G and P681H, and calculated the average to inform on the frequency of B.1.1.7 sequences in the sample.

### Statistical Analysis

Significance test were performed as indicated where appropriate. Statistical significance levels were determined as follows: ns = p > 0.05; * = p ≤ 0.05; ** = p ≤ 0.01; *** = p ≤ 0.001; **** = p ≤ 0.0001.

## Acknowledgements

We would like to thank Emmie de Wit, Brandi Williamson, Natalie Thornburg, Sue Tong, Sujatha Rashid, Ranjan Mukul, Kimberly Stemple, Craig Martens, Kent Barbian, Stacey Ricklefs, Sarah Anzick, Rose Perry, Tom Jones, Ryan Stehlik, Seth Cooley and Shanda Sarchette, and the animal care takers for their assistance during the study. The following reagent was obtained through BEI Resources, NIAID, NIH: SARS-Related Coronavirus 2, Isolate hCoV-19/England/204820464/2020, NR282 54000, contributed by Bassam Hallis.

## Author contributions

JRP, KCY, VJM designed the studies.

JRP, KCY, BJF, VA, MH, JES, NvD performed the experiments. JRP, KCY, CS analyzed results.

JRP, KCY, VJM wrote the manuscript. All co-authors reviewed the manuscript.

## Materials and Correspondence

All material requests should be sent to Vincent J. Munster.

## Supplementary Material

**Supplemental Figure 1:**
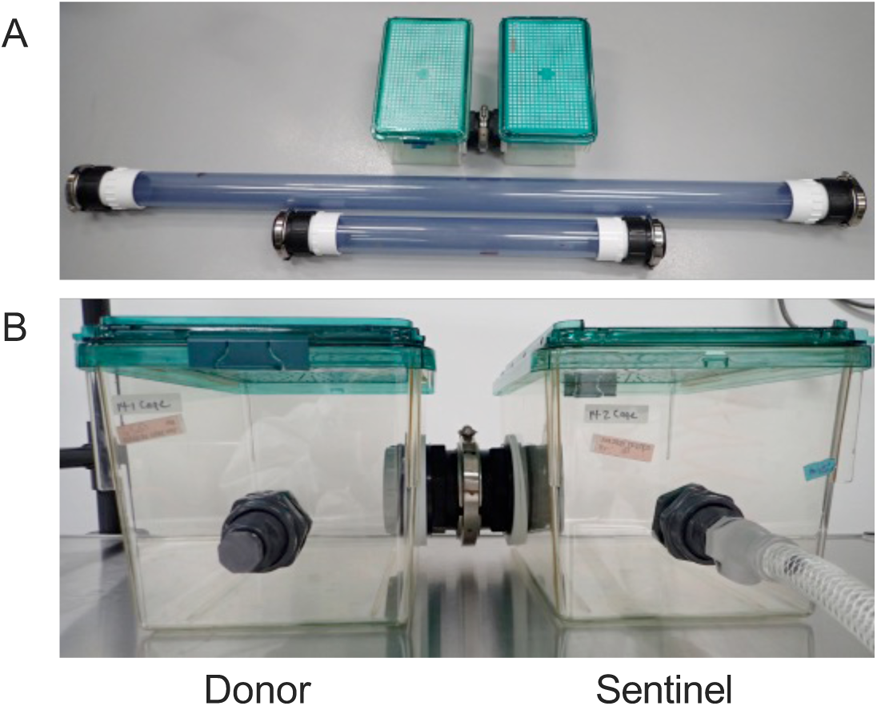
Aerosol transmission cages. **A/B.** Design of a new caging system in which two hamster cages could be separated at 3 different distances. The distances chosen were nominally 16.5 cm, 106 cm and 200 cm. The distance could be adapted by swapping out a 76 mm inside diameter connection tube. Cages were installed on autoclavable stainless steel shelves (Metro) inside a BSL-4 containment laboratory. Airflow was measured by flowmeters mounted to the shelves. Air was pulled through the system by a negative-pressure pump (Vacuubrand) and filtered through a hepa-filter before the exhaust.

**Supplementary Figure 2:**
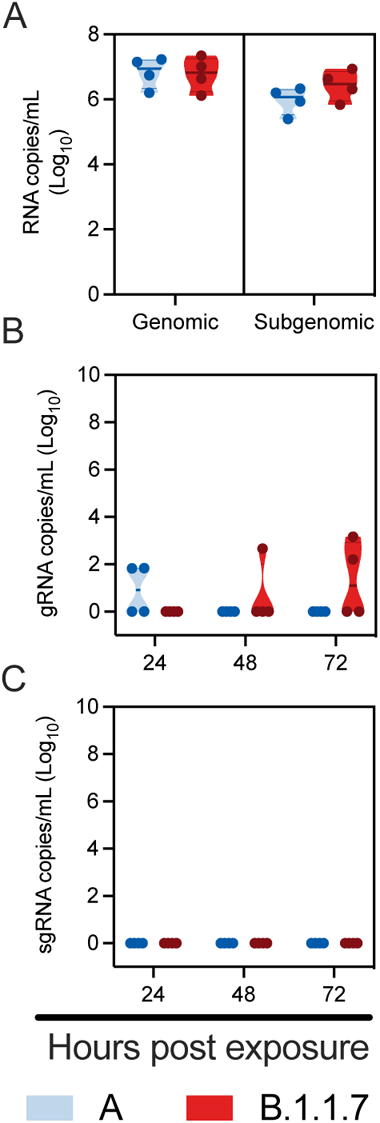
B.1.1.7 and lineage A aerosol transmission efficiency at three days post inoculation. Comparison of aerosol transmission efficiency of lineage A and B.1.1.7 SARS-CoV-2 variants in the Syrian hamster. Donor Syrian hamsters were inoculated with 8×10^4^ TCID_50_ SARS-CoV-2 lineage A or B.1.1.7 variant. After 72 hours, donors were introduced to the upstream cage and sentinels (2:2 ratio) into the downstream cage. Exposure was limited to one hour for B.1.1.7 and lineage A (N = 4, respectively). **A.** Viral load in oropharyngeal swabs of donors collected 72 hours post inoculation was measured by gRNA and sgRNA. **B/C.** To demonstrate transmission, sentinels were monitored for start and continuation of respiratory shedding. Viral load in oropharyngeal swabs of sentinels was measured by gRNA and sgRNA; swabs were collected at 24, 48 and 72 hours post exposure to the donors. Exposure at 200 cm distance. Truncated violin plots depicting median, quantiles and individuals, blue = lineage A, red = B.1.1.7. Abbreviations: A, lineage A variant; g, genomic; sg, subgenomic.

**Supplementary Figure 3:**
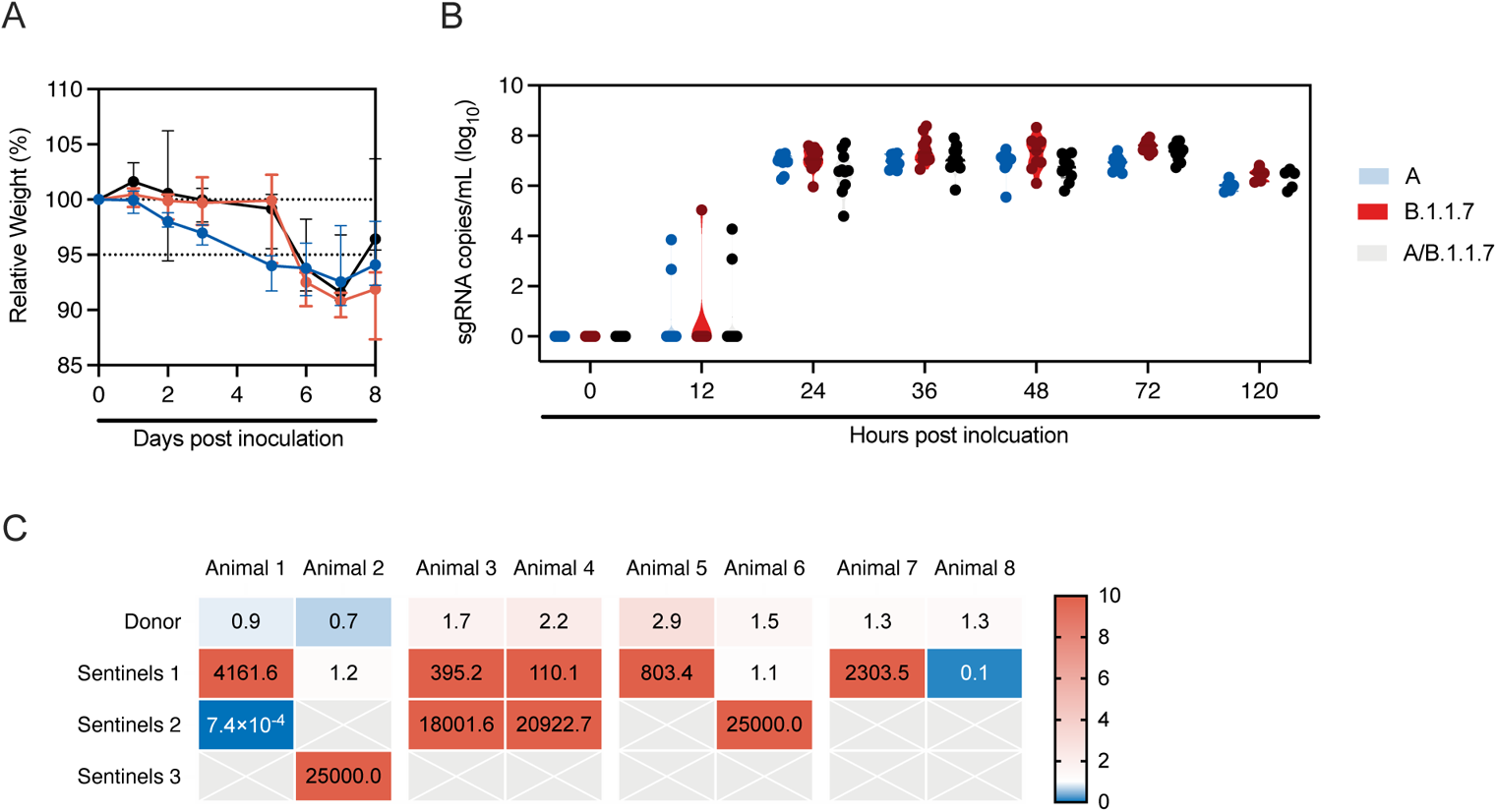
Dual infection with lineage A and B.1.1.7 variant in the Syrian hamster. Animals (N = 10) were inoculated with both lineage A and B.1.1.7 variant with 10^2^ TCID_50_ via the intranasal route (1:1 ratio), **A.** Relative weight loss in hamsters after dual inoculation in comparison to lineage A or B.1.1.7 variant inoculation. Graph shows median and 95% CI, N = 10. **B.** Respiratory shedding as measured by sgRNA in oropharyngeal swabs collected at 0, 12, 24, 36, 48, 72 and 120 hours post inoculation. Truncated violin plots depicting median, quantiles and individual, N = 10. **C.** Donor animals (N = 8) were inoculated with both lineage A and B.1.1.7 variant with 10^2^ TCID_50_ via the intranasal route (1:1 ratio), and three groups of sentinels (Sentinels 1, 2 and 3) were exposed subsequently at 16.5 cm distance. Ratio of B.1.1.7 and lineage A variant found in oropharyngeal swabs taken at day 2 post exposure/inoculation for each individual donor and sentinel, measured by duplex-qRT-PCR and depicted by ct foldchange (B.1.1.7 over lineage A variant). Colors refer to scale on the right. Samples for which only one variant was detected by PCR were set to 25,000. Abbreviations: A, lineage A variant; sg, subgenomic.

**Supplementary Table 1:** Aerosol Transmission Cage Validation Parameters. Special transmission cages were designed to model airborne transmission between Syrian hamsters. Volume of air (cages plus connection tube), air flow velocity in the tube and time for particles to traverse is provided at a cage air change rate of 30/h.

| Cage system | Volume (L) | Linear velocity (cm/min) | Time (sec) |
| --- | --- | --- | --- |
| Connecting tube (200 cm) | 1073 | 420 | 26.7692784 |
| Connecting tube (106 cm) | 945 | 370 | 15.059599 |
| Connecting tube (16.5 cm) | 837 | 327 | 2.80215197 |

**Supplementary Table 2:** anti-spike ELISA results for sentinels in aerosol transmission studies. Presence of SARS-CoV-2 spike IgG antibodies in sentinels co-housed at 16.5, 106 and 200 cm distance from lineage A or B.1.1.7(*) variant inoculated donor hamsters. Detected in serum obtained 14 days post exposure. negative (neg): optical density (at 450 nm) < 0.124, positive (pos) optical density (at 450 nm) ≥ 0.124.

| Condition | Animal | Seroconverted |
| --- | --- | --- |
| **16.5 cm** | S1 | pos |
|  | S2 | pos |
|  | S3 | pos |
|  | S4 | pos |
| **106 cm** | S1 | pos* |
|  | S2 | pos* |
|  | S3 | pos* |
|  | S4 | pos* |
|  | S5 | pos |
|  | S6 | pos |
|  | S7 | pos |
|  | S8 | pos |
| **200 cm** | S1 | pos* |
|  | S2 | pos* |
|  | S3 | pos* |
|  | S4 | pos* |
|  | S5 | pos |
|  | S6 | pos |
|  | S7 | pos |
|  | S8 | pos |

**Supplementary Table 3:**
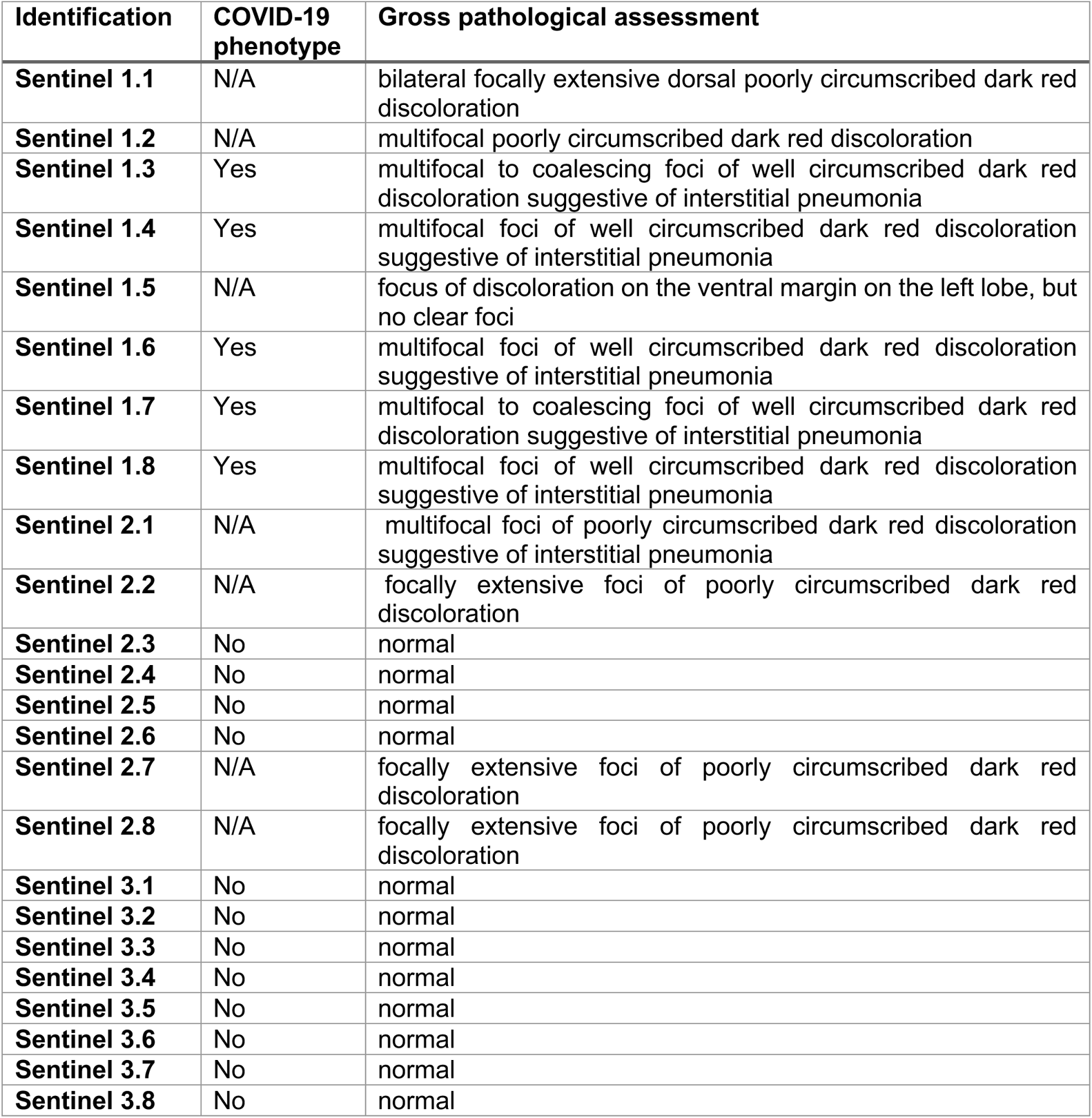
Pathological assessment of lungs collected at 5 days post exposure. Donor animals (N = 8) were inoculated with both lineage A and B.1.1.7 variant with 10^2^ TCID_50_ via the intranasal route (1:1 ratio), and three groups of sentinels (Sentinels 1, 2 and 3) were exposed subsequently at 16.5 cm distance. At five days post exposure, lungs were observed for gross pathology.

| Identification | COVID-19 phenotype | Gross pathological assessment |
| --- | --- | --- |
| **Sentinel 1.1** | N/A | bilateral focally extensive dorsal poorly circumscribed dark red discoloration |
| **Sentinel 1.2** | N/A | multifocal poorly circumscribed dark red discoloration |
| **Sentinel 1.3** | Yes | multifocal to coalescing foci of well circumscribed dark red discoloration suggestive of interstitial pneumonia |
| **Sentinel 1.4** | Yes | multifocal foci of well circumscribed dark red discoloration suggestive of interstitial pneumonia |
| **Sentinel 1.5** | N/A | focus of discoloration on the ventral margin on the left lobe, but no clear foci |
| **Sentinel 1.6** | Yes | multifocal foci of well circumscribed dark red discoloration suggestive of interstitial pneumonia |
| **Sentinel 1.7** | Yes | multifocal to coalescing foci of well circumscribed dark red discoloration suggestive of interstitial pneumonia |
| **Sentinel 1.8** | Yes | multifocal foci of well circumscribed dark red discoloration suggestive of interstitial pneumonia |
| **Sentinel 2.1** | N/A | multifocal foci of poorly circumscribed dark red discoloration suggestive of interstitial pneumonia |
| **Sentinel 2.2** | N/A | focally extensive foci of poorly circumscribed dark red discoloration |
| **Sentinel 2.3** | No | normal |
| **Sentinel 2.4** | No | normal |
| **Sentinel 2.5** | No | normal |
| **Sentinel 2.6** | No | normal |
| **Sentinel 2.7** | N/A | focally extensive foci of poorly circumscribed dark red discoloration |
| **Sentinel 2.8** | N/A | focally extensive foci of poorly circumscribed dark red discoloration |
| **Sentinel 3.1** | No | normal |
| **Sentinel 3.2** | No | normal |
| **Sentinel 3.3** | No | normal |
| **Sentinel 3.4** | No | normal |
| **Sentinel 3.5** | No | normal |
| **Sentinel 3.6** | No | normal |
| **Sentinel 3.7** | No | normal |
| **Sentinel 3.8** | No | normal |

**Supplementary Table 4:** qRT-PCR and sequencing results for donor and sentinel animals. Donor animals (N = 8) were inoculated with both lineage A and B.1.1.7 variant with 10^2^ TCID_50_ via the intranasal route (1:1 ratio), and three groups of sentinels (Sentinels 1, 2 and 3) were exposed subsequently at 16.5 cm distance. Viral load in copies/reaction (measured by qRT-PCR) and percentage of B.1.1.7 detected in oropharyngeal swabs taken at day 2 post exposure for each individual donor and sentinel, determined by deep sequencing (expressed as %) and expressed as fold-change over lineage A as measured by duplex-qRT-PCR.

| Animal | raw reads | qc reads | Viral RNA copies/reaction | % B.1.1.7 | PCR fold-change |
| --- | --- | --- | --- | --- | --- |
| Donor 1 | 9697 | 9289 | 1239 | 54 | 0.87408891 |
| Donor 2 | 77807 | 75255 | 85054 | 45 | 0.67126948 |
| Donor 3 | 78379 | 75218 | 103141 | 67.66667 | 1.69959284 |
| Donor 4 | 87992 | 85444 | 101401 | 70 | 2.1594464 |
| Donor 5 | 85062 | 82728 | 199462 | 74.66667 | 2.93509401 |
| Donor 6 | 135883 | 132657 | 254909 | 62 | 1.48700317 |
| Donor 7 | 45156 | 43638 | 71343 | 57.66667 | 1.2711726 |
| Donor 8 | 93687 | 91017 | 93356 | 57.66667 | 1.30263123 |
| Sentinel 1.1 | 65408 | 63416 | 126315 | 99 | 4161.61905 |
| Sentinel 1.2 | 41250 | 39943 | 38272 | 47 | 1.18602528 |
| Sentinel 1.3 | 112221 | 107500 | 20842 | 98.33333 | 395.151923 |
| Sentinel 1.4 | 107062 | 101891 | 62637 | 96 | 110.145084 |
| Sentinel 1.5 | 91154 | 88209 | 42500 | 96.66667 | 803.405938 |
| Sentinel 1.6 | 136060 | 131881 | 16994 | 58 | 1.12334412 |
| Sentinel 1.7 | 44594 | 43114 | 115352 | 98 | 2303.5488 |
| Sentinel 1.8 | 99311 | 96261 | 169096 | 16 | 0.12975316 |
| Sentinel 2.1 | 131545 | 126930 | 29513 | failed | 0.00073727 |
| Sentinel 2.3 | 33284 | 31942 | 4003 | 100 | 18001.5885 |
| Sentinel 2.4 | 25906 | 25000 | 4056 | 100 | 20922.7397 |
| Sentinel 2.6 | 10848 | 10219 | 522 | failed | * only B.1.1.7 detected |
| Sentinel 3.2 | 154787 | 141295 | 65 | failed | * only B.1.1.7 detected |

**Supplementary Table 5:**
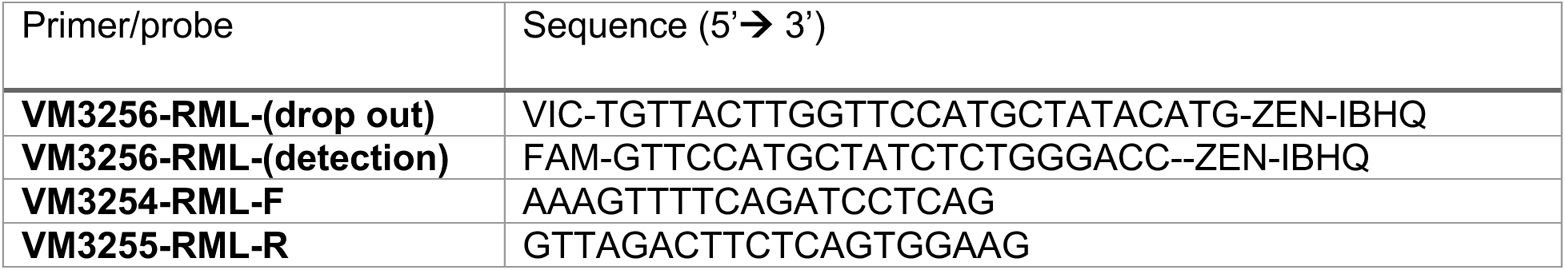
Duplex-qRT-PCR Primers and Probes.

**Supplementary Table 6:** Sequence results of virus stock B.1.1.7.

| ORF | a.a. change | Percentage |
| --- | --- | --- |
| nsp6 | D165G | 14 |
| nsp6 | L257F | 18 |
| nsp7 | V11I | 13 |

